# A positive feedback-based mechanism for constriction rate acceleration during cytokinesis in *C. elegans*

**DOI:** 10.1101/161133

**Authors:** Renat N. Khaliullin, Rebecca A. Green, Linda Z. Shi, J. Sebastian Gomez-Cavazos, Michael W. Berns, Arshad Desai, Karen Oegema

## Abstract

During cytokinesis, an equatorial actomyosin contractile ring constricts at a relatively constant overall rate despite its progressively decreasing size. Thus, the per-unit-length rate of ring closure increases as ring perimeter decreases. To understand this acceleration, we monitored cortical surface and ring component dynamics during the first division of the *C. elegans* embryo. We show that the polar cortex expands during ring constriction to provide the cortical surface area required for division. Polar expansion also allows ring myosin to compress cortical surface along the pole-to-pole axis, leading to a continuous flow of cortical surface into the ring. We propose that feedback between ring myosin and compression-driven cortical flow drives an exponential increase in the amount of ring myosin that maintains the high overall closure rate as ring perimeter decreases. We further show that an analytical mathematical formulation of the proposed feedback, called the Compression Feedback model, recapitulates the experimental observations.

**IMPACT STATEMENT:** During cytokinesis, positive feedback between myosin motors in the contractile ring and compression-driven cortical flow along the axis perpendicular to the ring drives constriction rate acceleration to ensure timely cell separation.

**MAJOR SUBJECT AREAS:** Cell biology, Computational and Systems Biology

## INTRODUCTION

During cytokinesis in animal cells, constriction of an equatorial actomyosin ring cinches the mother cell surface to generate a dumbbell-shaped structure with an intercellular bridge that connects the two daughter cells (Fededa and Gerlich, 2012; Green et al., 2012). Following chromosome segregation in anaphase, the contractile ring assembles in response to signaling by the anaphase spindle that activates RhoA at the cell equator (Green et al., 2012; Jordan and Canman, 2012; Piekny et al., 2005). RhoA patterns the equatorial cortex by recruiting contractile ring components from the cytoplasm (Vale et al., 2009; Yumura, 2001; Zhou and Wang, 2008). RhoA activates Rho kinase, which promotes the assembly and recruitment of myosin II (Matsumura et al., 2011) and the formin that assembles the long actin filaments that make up the ring (Otomo et al., 2005). Contractile rings also contain membrane-associated septin filaments (Bridges and Gladfelter, 2015) and the filament cross linker anillin (D'Avino, 2009; Piekny and Maddox, 2010). Recent work in the *C. elegans* embryo suggests that the equatorial cortex is compressed after this initial patterning, leading to the alignment of actin filament bundles as the ring forms (Reymann et al., 2016). After its assembly, the ring begins to constrict in the around-the-ring direction. Constriction is thought to be coupled to the progressive disassembly of the ring (i.e. loss of components in proportion to reduction in length) (Murrell et al., 2015; Schroeder, 1990).

Ring constriction must complete within a short cell cycle window during mitotic exit (Canman et al., 2000; Martineau et al., 1995; Straight et al., 2003). Timely constriction relies on the conserved ability of contractile rings to maintain a relatively constant overall closure rate despite their progressively decreasing perimeter (Biron et al., 2004; Bourdages et al., 2014; Calvert et al., 2011; Carvalho et al., 2009; Ma et al., 2012; Mabuchi, 1994; Pelham and Chang, 2002; Zumdieck et al., 2007). This property implies that the per-unit-length constriction rate increases as the rings get smaller. Prior work has suggested that this acceleration could arise if a constriction-rate controlling element is retained, rather than lost due to disassembly, as the ring shortens. For example, if myosin motors are not lost as the ring constricts, its concentration would increase in proportion to the reduction in perimeter, which could explain why the per-unit-length constriction rate increases as the ring shortens. Alternatively, it has been proposed that the number of actin filaments could be retained. If actin filaments shorten from their ends during constriction, the overall amount of actin polymer could decrease in proportion to the reduction in perimeter while the number of filament ends remains constant, perhaps leading to observed increase in the per-unit-length constriction rate (Carvalho et al., 2009).

Here, we explore the mechanisms underlying constriction rate acceleration during the first division of the *C. elegans* embryo. By generating a 4D map of cortical surface dynamics, we show that cortex at the cell poles expands in response to the tension generated by the constricting ring to provide the increased cortical surface area required to generate the daughter cells. The ability of the polar cortex to expand in response to tension also allows ring myosin to compress cortical surface along the pole-to-pole axis perpendicular to the ring, leading to a continuous flow of cortical surface into the ring during constriction. We show that the ring compresses cortical surface throughout cytokinesis at a rate proportional to the amount of ring myosin. In addition, the amount of ring myosin increases in proportion to the amount of cortical surface pulled into the ring by compression. The per-unit-length amount of ring myosin and the per-unit-length rates of cortical compression and ring constriction increase with the same exponential kinetics as the ring closes, suggesting control by positive feedback. Based on our observations, we propose that feedback between ring myosin and compression-driven cortical flow drives ring myosin accumulation, which in turn increases the per-unit-length constriction rate to keep the overall constriction rate high as the ring closes. We show that an analytical mathematical formulation of the proposed feedback, called the Compression Feedback model, recapitulates our experimental observations.

## RESULTS

### The cortex at the cell poles expands in response to tension generated by the constricting ring without limiting the constriction rate

During the first division of the *C. elegans* embryo, the surface area of the cell increases by ~40% to accommodate the shape change that generates the daughter cells. Work in multiple systems has shown that the entire cell surface, from cortex-associated granules in the cytoplasm to cell surface receptors, moves in a coordinated fashion during cytokinesis (Cao and Wang, 1990; Dan, 1954; Dan and Dan, 1940; Dan et al., 1938; DeBiasio et al., 1996; Fishkind et al., 1996; Hird and White, 1993; Reymann et al., 2016; Swann and Mitchison, 1958; Wang et al., 1994). In a classic set of experiments, Dan and colleagues measured the distance between surface adhered particles to monitor changes in cortical surface area (compression and expansion) during cytokinesis in sea urchin embryos. This analysis revealed that ring constriction occurs coincident with a wave of cortical expansion that initiates at the cell poles and propagates towards the furrow (Dan et al., 1938; Dan and Ono, 1954; Dan et al., 1937; Swann and Mitchison, 1958). Although these experiments provided a rough map of where expansion occurs, they did not allow quantification of the extent of change in cortical surface area or provide a map of cortical surface movements. Note that the analysis of cell surface dynamics described above refers to movement, expansion and compression of the cortex and associated structures. How deposition of plasma membrane, the fluid lipid layer that overlies the cortex, is controlled and where it occurs are distinct questions that we will not discuss here.

To generate a quantitative map of cortical surface dynamics during the first division of the *C. elegans* embryo, we employed an updated version of the classical approach in which we used myosin foci rather than surface adhered particles as fiduciary marks. We imaged the cortex at high time resolution (2s intervals, cyan box in Figure 1A, Video 1) in embryos expressing a GFP fusion with the heavy chain of non-muscle myosin II (NMY-2; hereafter myosin::GFP; Figure 1 - Figure Supplement 1A,B). In addition to its RhoA-dependent enrichment in the contractile ring, myosin is in small puncta, distributed over the entire cortex, that flow together with actin filaments (LifeAct::mKate2, Figure 1 - Figure Supplement 1C), validating their utility as fiduciary marks for monitoring cortical movements. To temporally and spatially align data collected in different embryos, ring constriction was also monitored at lower time resolution in the same embryos (36s intervals, Figure 1A, Figure 1 - Figure Supplement 2). Because the contractile ring closes asymmetrically within the division plane ((Maddox et al., 2007); Figure 1A, Figure 1 - Figure Supplement 2), cortical dynamics are not cylindrically symmetric. Therefore, we generated an average 4D map of cortical movement by computationally combining data from 93 embryos imaged in random rotational orientations (Figure 1A, Figure 1 - Figure Supplement 2). We defined the top of the embryo as the side where the furrow ingresses first, the bottom as the opposite side, and referenced positions around the embryo circumference by the angle θ relative to the initial ingression axis (Figure 1A). For temporal alignment, we fit a line to normalized ring size 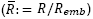 versus time between 30% and 80% closure for each embryo, and extrapolated this line to 1 and 0 to define t_0_ (cytokinesis onset) and t_CK_ (time of cytokinesis), respectively (Figure 1A, Figure 1 - Figure Supplement 2). Cortical movement could not be monitored in the division plane, because it is hidden inside the cell, or at the cell poles, due to their high curvature. Thus, this approach provided a quantitative picture of cortical movement in the central 2/3 of the embryo throughout cytokinesis (Figure 1B; Video 2).

**Figure 1.**
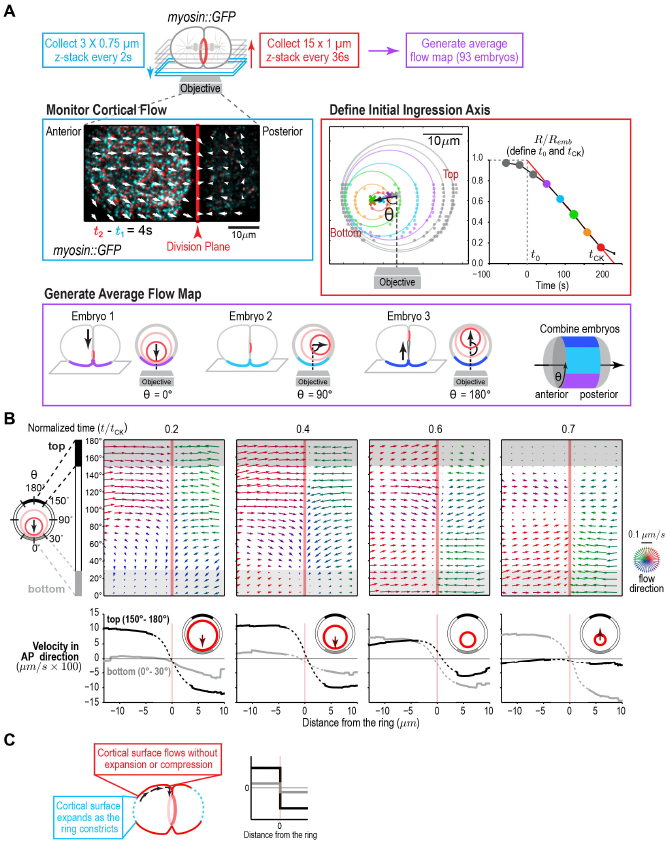
A quantitative map of cortical surface dynamics during the first cytokinesis in the *C. elegans* embryo reveals that the cortical surface at the cell poles expands as the ring constricts. (**A**) *(top)* Schematic of the experimental procedure. *(middle, left)* Superposition of images of the cortex acquired 4s apart. Arrows indicate cortical flow (magnified 2.5X). *(middle, right)* The initial ingression axis, t_0_, and t_CK_ were defined as shown for a representative embryo. The angle 0 specifies the position of the imaged cortex relative to the initial ingression axis. Image and quantification are representative of the 93 imaged embryos. *(bottom)* Angular position was used to combine data from 93 embryos to generate an average flow map. (**B**) *(top)* Average flow at the indicated timepoints. Arrows show direction and magnitude of the displacement in 1s (magnified 20X). *(middle)* Graphs are average velocity in the A-P direction versus position along the A-P axis for the cortex on the top *(black)* and bottom (grey) of the embryo *(shaded in flow maps).* Surface movement changes direction across the division plane, the apparent velocity gradient close to the division plane is a projection artifact due to the fact that the cortical surface turns inwards as it approaches the furrow from either side (dotted regions on velocity curves). (**C**) Schematics show the predicted cortical velocity profile along the AP axis if surface is gained at the poles; velocity would be constant in magnitude within the flow map region with opposite directions on the two sides of the ring, as is experimentally observed.

The 4D map allowed us to determine where cortical surface expansion occurs as the ring closes in the *C. elegans* embryo. Prior work monitoring the movement of surface adhered particles in sea urchin and *Xenopus* embryos indicated that surface expansion occurs at the poles and immediately behind the contractile ring, respectively, in these systems (Bluemink and de Laat, 1973; Byers and Armstrong, 1986; Danilchik et al., 2003; Gudejko et al., 2012; Selman and Perry, 1970; Swann and Mitchison, 1958). In addition to these two patterns, we also considered the possibility that the cortex would expand uniformly, an assumption often used in models of cytokinesis (Turlier et al., 2014; Zumdieck et al., 2007). Each of these three patterns predicts a different profile for the Anterior-Posterior (AP) component of cortical velocity along the embryo. For uniform surface expansion, a gradient of velocities is predicted, where the cortical velocity immediately behind the ring equals the velocity of furrow ingression and the velocity decreases linearly towards the cell poles. For surface expansion behind the ring, no cortical movement is predicted on the observable embryo surface. If surface expansion is limited to the poles, the cortical velocity is predicted to be constant within the flow map region (Figure 1 - Figure Supplement 3). The cortical velocity profile measured from the flow map indicated that the cortical surface at the cell poles expands as the ring constricts, whereas the cortex between the poles and the division plane flows at constant velocity towards the division plane, without expansion or compression (Figure 1B). Note that the apparent velocity gradient that spans the division plane (Figure 1B, *dashed regions on velocity curves)* is a projection artifact due to the fact that the cortical surface turns inwards as it approaches the furrow from either side. As expected, based on the asymmetric closure of the contractile ring within the division plane, the velocity of cortical flow was higher on the top of the embryo during the first half of cytokinesis when the furrow ingresses from the top (Figure 1B, black traces) and became higher on the bottom of the embryo towards the end when the furrow ingresses from the bottom (Figure 1B, grey traces; Video 2).

Cutting the cortex parallel to the division plane using a laser revealed that the cortex is under tension during cytokinesis (Figure 2A). To determine if cortical tension limits the constriction rate, we assayed the effect of the cortical cuts on ring closure. Cortical cuts spanning the visible area of cortex on the anterior side of the embryo (~10um in length) were made parallel to the division plane when the ring was at ~50% closure, and the effect on contractile ring closure rate was assessed by measuring the difference in ring sizes immediately before and 13s after the cut. The cortical opening resulting from the ablation was approximately 35μm^2^, which would be expected to increase the constriction rate from the control rate of 0.22 + 0.5 μm/s to ~0.25 μm/s over our 13s interval if the cortical surface tension is the dominant force limiting the ring closure rate (**see Methods for details**). In contrast, the measured constriction rate after was not increased after cutting (0.18 + 0.03 μm/s; Figure 2B,C), indicating that cortical tension does not impose significant resistance to ring pulling. Cuts made perpendicular to the ring also had no effect on the constriction rate (0.19 + 0.03 μm/s **data not shown**). Consistent with the results of the laser cutting experiments, inhibiting the Arp2/3 complex by depleting its ARX-2 subunit, which is expected to reduce effective cortical viscosity and thus cortical tension (Chaudhuri et al., 2007; Davies et al., 2014; Tseng and Wirtz, 2004), also did not alter the constriction rate (Figure 2 - Figure Supplement 1A).

**Figure 2.**
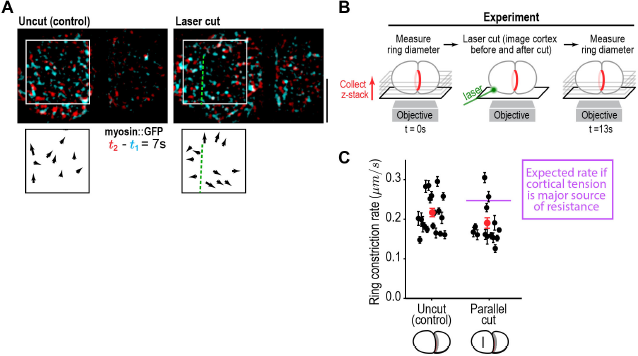
Cortical tension does not limit the rate of ring closure. (**A**) The success of cortical cuts was assessed by comparing surface images of cortical myosin before *(cyan)* and after *(red)* the cut to monitor the movement of myosin foci away from the cut site. Representative images are shown. Scale bar is 10 μm. (**B**) Schematic of laser ablation experiment to determine if cortical resistance limits the rate of contractile ring closure. Contractile ring sizes were measured from z-stacks acquired before and 13s after a cut was made across the cortex with a laser. (**C**) Graph plots the rates of ring closure derived from before and after ring size measurements for uncut controls (n=19 embryos) and embryos with cuts parallel to the division plane (n=14 embryos). Black symbols are single embryo measurements with measurement errors. Red symbols are the means; error bars are the SEM. The purple line marks expected closure rate if cortical tension is a major source of resistance.

Putting the results of our flow map analysis with our laser cutting and Arp2/3 inhibition experiments together, we conclude that the cortex at the poles expands in response to tension generated by the constricting ring without providing significant resistance that would affect the rate of ring closure. In contrast, the cortex in the region between the ring and the poles flows towards the ring without expansion or compression. The differential response of the polar cortex to ring-generated tension is consistent with the idea of polar relaxation hypothesized in early conceptual models of cytokinesis (Greenspan, 1978; Swann and Mitchison, 1958; Taber, 1995; White and Borisy, 1983; Wolpert, 1960; Zinemanas and Nir, 1987; Zinemanas and Nir, 1988), and suggests that the polar cortex has unique mechanical properties compared to the intervening cortex that does not expand (see discussion for possibilities). The fact that cortical tension does not limit the rate of ring constriction suggests that the constriction rate is instead limited by ring internal friction. We conclude that the viscosity of the polar cortex is negligible compared to the viscosity internal to the ring; thus, ring myosin generated force primarily counters ring internal friction to drive ring constriction (Figure 2 - Figure Supplement 1B). Ring constriction, in turn, affects cortical tension and drives expansion of polar cortex.

### Ring myosin compresses cortical surface along the pole-to-pole axis perpendicular to the ring, pulling in new cortical surface at a rate proportional to the amount of ring myosin

In the *C. elegans* embryo, as in other systems, spindle-based signaling activates RhoA on the equatorial cortex following anaphase onset leading to the recruitment of contractile ring proteins including myosin II, the septins, and anillin (Jenkins et al., 2006; Maddox et al., 2005; Maddox et al., 2007; Mangal et al., 2018; Motegi and Sugimoto, 2006; Schonegg et al., 2007; Tse et al., 2012; Werner et al., 2007). An astral microtubule based mechanism that clears contractile ring proteins from the polar cortex also confines contractile ring protein recruitment to a defined equatorial zone (Mangal et al., 2018; Werner et al., 2007). Prior work in the *C. elegans* embryo has suggested that the equatorial cortex is compressed during contractile ring assembly, coincident with the alignment of actin filament bundles to form the ring (Reymann et al., 2016). Cortical surface compression is detected as a gradient in the velocity of cortical surface flow. Consistent with the idea that cortical surface is compressed during contractile ring assembly, we observed a linear gradient in the velocity of cortical flow that spanned the cell equator in our flow map at early time points prior to furrow ingression (Figure 3A). The linear gradient indicated that, during contractile ring assembly when the ring is on the embryo surface, cortical surface is uniformly compressed across a 10 μm wide region along the perpendicular-to-the-ring axis between the two relaxing poles.

**Figure 3.**
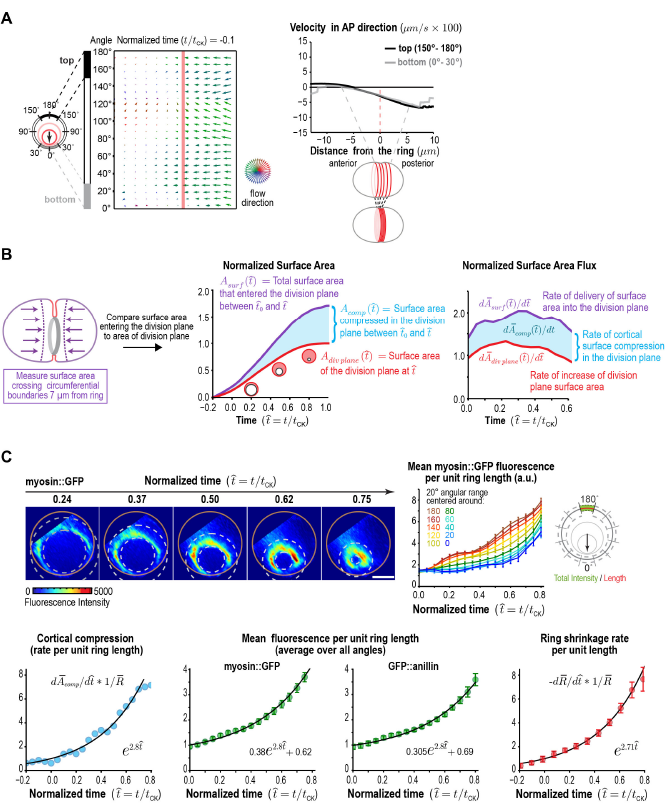
Ring myosin compresses cortical surface along the axis perpendicular to the ring, pulling in new cortical surface at a rate proportional to the amount of ring myosin. (**A**) The equatorial cortex is compressed during contractile ring assembly. Following the onset of spindle-based RhoA signaling, the initial recruitment of contractile ring proteins leads to uniform compression of cortical surface along the axis perpendicular to the forming ring across a 10 pm wide region spanning the cell equator. (*left)* Average flow map at (t/t_CK_=-0.1) immediately after the onset of spindle-based signaling (n= 93 embryos). *(middle)* The surface velocity profile reveals a linear velocity gradient that spans the cell equator (-5 to +5 pm), indicating a zone of cortical compression. (B) Cortical compression within the ring continues during constriction. *(left graph)* Plot comparing the area of the forming division plane *(red)* with the total cortical surface area that entered the division plane from the start of cytokinesis *(purple;* calculated as indicated in the schematic). *(right graph)* Plot comparing the rate of delivery of cortical surface into the division plane *(purple)* with the rate of growth of the division plane (red). The difference between the two is the rate of cortical surface compression *(cyan).* (**C**) The per-unit-length amount of ring myosin and the rate of cortical compression increase with the same exponential kinetics. *(top left)* Representative images of the division plane in embryos expressing myosin::GFP reconstructed from 40- plane z-stacks. Gold circles mark the embryo boundary and dashed circles mark the boundaries used for ring intensity measurements. *(top right)* Graph plots per-unit-length myosin::GFP fluorescence for the indicated angular ranges (n=36 embryos). *(bottom left)* Graph plots the rate of cortical surface compression per unit ring length (n=93 embryos). *(bottom middle)* Graphs plot mean per-unit-length myosin::GFP (n=36 embryos) and GFP::anillin (n=26 embryos) fluorescence (n=36 embryos) in the ring. *(bottom right)* Graph plots the per-unit-length rate of ring closure. Black lines are fitted single exponentials. Error bars are the SEM.

After its assembly, the ring begins to constrict in the around-the-ring direction, which has been proposed to be coupled to the progressive disassembly of the ring (i.e. loss of components in proportion to reduction in length) (Murrell et al., 2015; Schroeder, 1990). During constriction, the ring pulls the cortex behind it, which leads to a flow of cortex into the division plane. We were interested in whether the compression along the perpendicular-to-the-ring axis is limited to contractile ring assembly, or whether it might also continue during ring constriction. If compression stops, the constricting ring would generate the division plane by pulling the cortex behind it, and the cortical surface area entering the division plane would equal the area of the division plane. In contrast, if compression along the axis perpendicular to the ring continues during constriction, the cortical surface area entering the division plane would be larger than the area of the division plane by the amount of surface compressed.

To distinguish between these possibilities, we used the 4D cortical flow map to measure the cortical surface area entering the division plane and compare it to the area of the division plane (accounting for the fact that two surfaces are *generated-red outline in* Figure 3B). This analysis revealed that the area of the cortical surface that entered the division plane during ring constriction was significantly greater than the area of the division plane (Figure 3B, *middle panel).* The flux of cortical area into the division plane was 1.5 to 2-fold higher than the rate of change in the area of the division plane throughout cytokinesis, indicating ongoing cortical surface compression (Figure 3B, *right panel).* In control embryos, more cortex flowed in from the posterior side than from the anterior side, likely due to distinct mechanical cortical properties that arise downstream of the polarity machinery. Prior work showed that Arp2/3 inhibition impairs the recruitment of PAR-2 to the posterior cortex and makes myosin and actin dynamics on the posterior cortex more similar to those in embryo anterior (Xiong et al., 2011). Inhibiting the Arp2/3 complex by depleting ARX-2 abolished the difference between the two sides, but did not change the difference between the total amount of cortex entering the division plane and the area of the plane (Figure 3 - Figure Supplement 1; Video 3). This result suggests that the compression of cortical surface along the axis perpendicular to the ring persists throughout constriction, resulting in a continuous flow of cortical surface into the ring.

Next, we probed the relationship between the rate of cortical surface area compression along the axis perpendicular to the ring and the levels of two contractile ring components, myosin, which is required for ring constriction and cortical surface compression (Reymann et al., 2016; Shelton et al., 1999), and anillin, a filament cross-linker that localizes to the ring but is not essential for constriction or compression (Maddox et al., 2005; Maddox et al., 2007; Reymann et al., 2016). To do this, we monitored *in situ-tagged* myosin::GFP (Dickinson et al., 2013) (Figure 3C) and GFP::anillin (Figure 3 - Figure Supplement 2) in end-on reconstructions of the division plane. Both ring components exhibited similar behavior. Because overall measurements of ring component levels and constriction/compression rates scale with ring size, all of our analysis considers measurements per unit of ring length, which capture the evolution of the material properties of the ring independent of size. Quantification of mean per-unit-length fluorescence around the ring (after attenuation correction; Figure 3 - Figure Supplement 3) revealed a steady increase for both markers as constriction proceeded. The increase in the per-unit-length amounts of myosin and anillin began on the top of the ring, which ingresses first, and initiated later on the bottom, which ingresses after the constriction midpoint (Figure 3C, Figure 3 - Figure Supplement 2). Comparing the per-unit-length rate of cortical compression along the axis perpendicular to the ring to the per-unit-length amounts of myosin and anillin revealed that both increased with the same exponential kinetics during constriction (Figure 3C). Thus, new cortical surface is pulled into the ring due to cortical compression at a rate proportional to the amount of ring myosin. Like the rate of cortical compression along the axis perpendicular to the ring, the per-unit-length constriction rate also increased in proportion to the per-unit-length amount of myosin (Figure 3C). The exponential increase in the per-unit-length constriction rate explains the observed ability of the contractile ring to close at a relatively constant rate despite its progressively decreasing perimeter (Bourdages et al., 2014; Carvalho et al., 2009; Zumdieck et al., 2007). A relatively constant overall rate of ring closure is observed over a significant portion of constriction (Figure 1A; *t* = 50-200s) because the exponential increase in the constriction rate balances the decrease in ring size.

We note that in prior work in 4-cell stage *C. elegans* embryos, we had shown that myosin, anillin and septins levels in the ring increase ~1.3-fold as ring perimeter decreases 2-fold (from 50 to 25 μm), but had not concluded that contractile ring component accumulation was exponential. This is because the range of ring sizes between furrow formation and contact with the midzone, which occurs at a perimeter of ~25 μm and alters ring properties (Carvalho et al., 2009), is much smaller at the 4-cell stage than at the 1-cell stage. Although not sufficient to demonstrate exponential accumulation on their own, the 4-cell data are well fit by the same exponential equation that describes myosin and anillin accumulation at the 1-cell stage (Figure 3 - Figure Supplement 4), suggesting that ring components accumulate in a similar fashion across the first four cell divisions in the *C. elegans* embryo.

### An analytical mathematical model for positive feedback-mediated evolution of the contractile ring: the Compression Feedback model

From our experimental work we conclude that: (1) the ring compresses cortical surface along the axis perpendicular to ring constriction throughout cytokinesis at a rate proportional to the amount of ring myosin and, (2) the amount of ring myosin and anillin increase at a rate proportional to the rate at which cortical surface is compressed into the ring, (3) the per-unit-length amounts of ring myosin and anillin and the per-unit-length rates of cortical compression and ring constriction increase with the same exponential kinetics as the ring closes. The fact that ring components accumulate with exponential kinetics further suggests control by positive feedback. Our results suggest that the relevant feedback could be between the amount of ring myosin and the rate of cortical surface compression within the ring (i.e. ring myosin would lead to cortical compression that would deliver myosin into the ring). To explore this idea, we developed an analytical mathematical formulation, which we call the Compression Feedback model, consisting of three equations with three model parameters, that describes this feedback and can recapitulate our experimental results (Figure 4A,B).

**Figure 4.**
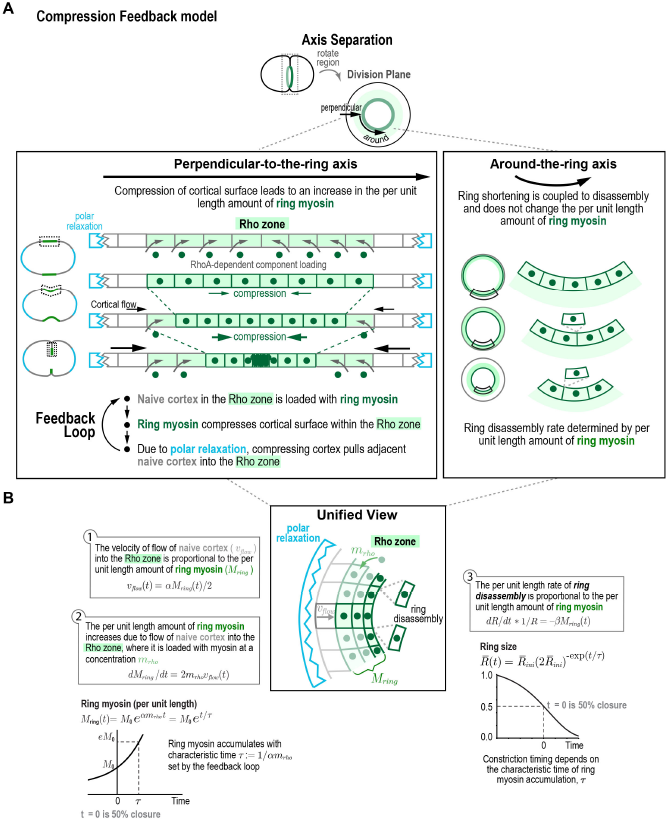
Compression Feedback model of cytokinesis. (**A**) The natural coordinate system for contractile ring dynamics has two axes, an axis parallel to ring constriction *(around-the-ring axis)* and an axis perpendicular to the ring *(perpendicular-to-the-ring axis).* Polar relaxation and filament alignment in the around-the-ring direction lead to anisotropy in behavior along the two axes, which are illustrated separately here. Along the axis perpendicular to the ring, feedback between ring myosin and compression-driven cortical flow leads to an exponential increase in the per-unit-length amount of ring myosin. Along the around-the-ring axis, constriction is coupled to disassembly and does not change the per-unit-length amount of ring myosin. (**B**) Formulation of the proposed mechanisms as an analytical mathematical model consisting of three equations and three model parameters. *(left)* Equations (1) and (2) describe the feedback loop between the amount of ring myosin and the velocity of compression-driven flow of cortical surface into the ring. Solving these equations gives the expression for the per-unit-length amount of ring myosin, which accumulates exponentially as shown in the graph. *(right)* The feedback loop operating perpendicular to the ring controls the per-unit-length amount of ring myosin, which in turn controls the per-unit-length rate of ring constriction as described in equation (3). Graph plots the equation for ring size resulting from solving the model equations in the time reference where *t* = 0 is the halfway point of ring closure.

The natural coordinate system for contractile ring dynamics has two axes, an axis parallel to ring constriction (Figure 4A, *around-the-ring axis)* and an axis perpendicular to the ring (Figure 4A, *perpendicular-to-the-ring axis).* Our experimental results suggest that polar relaxation leads to differential behavior in these two directions. After anaphase onset, spindle based signaling patterns the cortex, generating an equatorial zone, that has been termed the Rho zone (Bement et al., 2006; Green et al., 2012; Jordan and Canman, 2012; Piekny et al., 2005), where RhoA promotes the recruitment of contractile ring components including myosin and anillin (Jenkins et al., 2006; Maddox et al., 2005; Maddox et al., 2007; Mangal et al., 2018; Motegi and Sugimoto, 2006; Schonegg et al., 2007; Tse et al., 2012; Werner et al., 2007). The Rho zone occupies the central region of the pole-to-pole axis perpendicular to the ring. As our data indicate (Figure 3A), the initial recruitment of contractile ring proteins after anaphase onset results in uniform cortical compression across this central 10 μm wide zone. We propose that, due to polar relaxation, the compressing cortex pulls naïve cortex not patterned by the initial round of RhoA signaling, into the Rho zone (Figure 4A). The new cortex that flows into the Rho zone as a result of compression would be loaded with contractile ring components that would initiate compression and contribute to compression-driven cortical flow. Thus, *along the perpendicular-to-the-ring* axis a feedback loop would operate in which myosin in the ring compresses cortical surface, which pulls more surface that is loaded with myosin into the ring (Figure 4A, left panel). *In the around-the-ring direction,* reduction in ring perimeter would be coupled to disassembly (loss of ring components in proportion to reduction in length), with the per unit length rate of ring disassembly being determined by the per unit length amount of myosin. Thus, unlike the feedback loop operating along the perpendicular-to-the-ring axis, which would lead to an exponential increase in the per-unit-length levels of ring components, ring shortening would be coupled to disassembly and would not alter the per-unit-length amount of ring components.

In the mathematical formulation (Figure 4B), naïve cortex flows into the Rho zone at a velocity (*V*_flow_(*t*)) proportional to the per-unit-length amount of ring myosin (*M_ring_(t*); Figure 4B, Eqn. (1)), with *α* being the proportionality constant that relates the two. Ring myosin, in turn, increases at a rate proportional to this flow and the concentration of myosin that is loaded onto the cortex when it enters the rho zone *(m_rh0_*; Figure 4B, Eqn. (2)). As a result of the positive feedback between ring myosin and compression-driven flow, ring myosin increases exponentially with a characteristic time *τ := 1/αm_rh0_* (time required for ring myosin to increase ~2.7 fold; Figure 4B, bottom graph). The per-unit-length rate of ring constriction 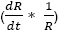 is proportional to the per-unit-length amount of ring myosin, related by the proportionality constant *β* (Figure 4B, Eqn. (3)). To avoid the difficulty of accurately assigning the exact point when cytokinesis starts, we solved these equations in the time reference where *t* = 0 is the halfway point of ring closure 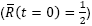. In this time reference, the equation for ring size is:

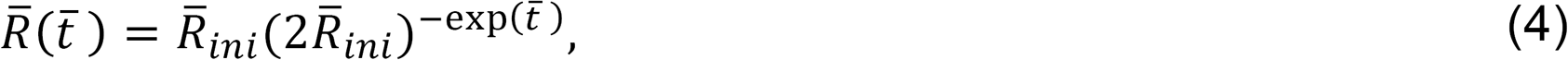

where 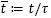 and 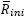 is the dimensionless characteristic ring size (held fixed at a value of 1.1; see Methods; Figure 4B, *right graph).* Other components, like anillin, that localize to the cell cortex will be delivered to the contractile ring via the same process as myosin, and would accumulate in a similar fashion, with

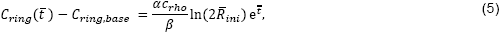

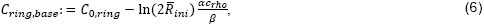

where *C_0,ring_* is the per-unit-length amount of the component at the half-way point of ring closure, *C_ring,base_* is the baseline amount of the ring component that does not increase exponentially, and *c_rh0_ (m_rh0_* for myosin) is the concentration of the component loaded onto naïve cortex when it enters the rho zone. The velocity of cortical flow and the constriction rate are

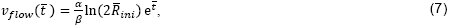

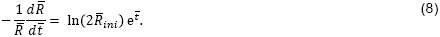

Thus, the per-unit-length constriction rate, velocity of cortical flow, and ring component amounts would all increase exponentially with the characteristic time of ring myosin accumulation (τ = *1/αm_rh0_*) set by the feedback loop between ring myosin and cortical flow, as we have observed experimentally (Figure 3C). We conclude that an analytical mathematical formulation that describes a feedback loop between ring myosin and compression-driven cortical flow can recapitulate the experimentally observed pattern of cortical surface compression and ring component and constriction dynamics.

### Fluorescence recovery after photobleaching of the division plane is consistent with the Compression Feedback model

The Compression Feedback model is characterized by anisotropy in the behavior in the perpendicular-to-the-ring and around-the-ring directions (Figure 5A). Along the perpendicular-to-the-ring direction, cortical compression within the ring pulls in cortical surface, which increases the per-unit-length amount of ring components and, as a consequence, the per-unit-length constriction rate. In contrast, in the around-the-ring direction, constriction is coupled to disassembly and does not affect the per-unit-length amount of ring components. An alternative model that could explain the increase in the per-unit-length amount of ring components, which we refer to as “Retention” model, is that the per-unit-length constriction rate accelerates due to retention of myosin and/or other ring components during ring shortening (Figure 5A). In the Retention model, compression in the around-the-ring direction increases the per-unit-length amount of ring components. In this model, myosin and anillin would not be lost due to disassembly, and their total amounts in the ring would remain constant during constriction, resulting in an increase in their per-unit-length amounts in inverse proportion to the reduction in ring size (levels would increase as 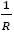). In the perpendicular-to-the-ring direction, compression would still pull cortical surface into the ring, as we have shown occurs experimentally, but the Retention model assumes that flow would not deliver myosin into the ring, either because levels of myosin on the delivered cortex are insignificant relative to the amount of myosin in the ring or because the delivered myosin is lost due to disassembly. Comparison with the total amounts of ring myosin and anillin suggested that, whereas the Retention model fit the data well for *t/t_ck_* between 0.2 and 0.6, there was significant deviation for timepoints outside of this range. In contrast, the Compression Feedback model fit the data well over the entire measured interval *(t/t_ck_* = 0.0 to 0.8; Figure 5B, Figure 5—Figure Supplement 1). The exponential accumulation predicted by the Compression Feedback model also fit the experimental data for the per-unit-length rates of ring shrinkage and cortical compression significantly better than the Retention model, which would predict that these rates, like the amount of ring myosin, would also increase as 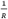 (Figure 5B).

**Figure 5.**
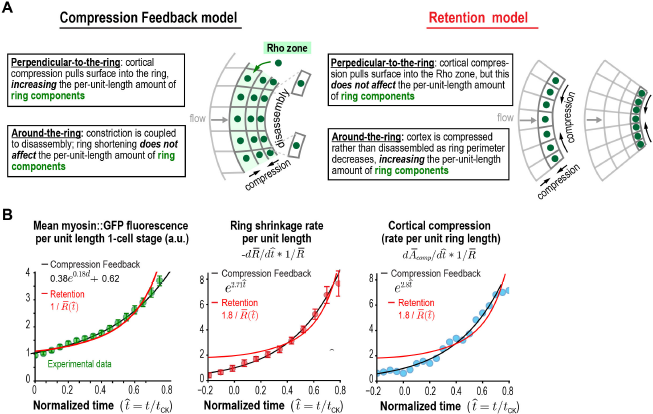
Myosin and anillin accumulation and the rates of ring constriction and cortical compression support the Compression Feedback model.

(**A**) Two models could explain the acceleration in the per-unit-length constriction rate during constriction. In the Compression Feedback model, acceleration results from a feedback loop between ring myosin and compression-driven cortical flow in the perpendicular-to-the-ring direction. In the Retention model, compression without disassembly in the around-the-ring direction increases the per-unit-length amounts of ring components. (**B**) Graphs show mean per-unit-length myosin::GFP fluorescence in the ring along with the per-unit-length constriction and cortical compression rates. Myosin fluorescence data is reproduced from Figure 3C to allow comparison of the best fits for the Compression Feedback *(black lines)* and Retention *(red lines)* models.

As a further test of the Retention and Compression Feedback models, we photobleached myosin in the entire division plane at ~30% closure, and monitored its subsequent recovery in the ring (Figure 6A). In prior work, we photobleached three contractile ring components (myosin::GFP, GFP::anillin, and GFP::septin) in contractile rings at the 4-cell stage. This analysis suggested that in contrast to myosin on the cortex outside of the ring, which has been shown to turn over rapidly (t_1/2_ of ~30s; (Mayer et al., 2010; Salbreux et al., 2012)), significant turnover due to exchange with components in the cytoplasm was not observed for myosin, anillin, or the septins in the ring. The Retention model predicts that cortical compression along the perpendicular-to-the-ring direction does not contribute to ring myosin accumulation. In the around-the-ring direction, the cortex is compressed as ring perimeter decreases leading to an increase in the per-unit-length amount of both the bleached and residual fluorescent myosin in proportion to the reduction in ring perimeter (both would increase as 1/R; (Figure 6B, *top).* The Compression Feedback model predicts that after the myosin in the ring is bleached, cortical compression in the perpendicular-to-the-ring direction will continue to pull continue naive cortex into the Rho zone that will be loaded with fluorescent myosin from the cytoplasm. Thus, after the bleach, the per-unit-length amount of fluorescent myosin in the ring will rapidly begin to increase again at an exponential rate comparable to that in controls. In the around-the-ring direction, the bleached myosin in the ring will be disassembled in proportion to the reduction in ring length; thus, the per-unit-length amount of bleached myosin in the ring will remain constant (Figure 6B, *bottom).* Our data indicated that the per-unit-length amount of fluorescent myosin in the ring increased exponentially following bleaching at a rate comparable to that in controls, and the difference between the fluorescence in the control and bleached embryos, which is the amount of bleached myosin, remained constant as the ring constricted. These observations are consistent with the predictions of Compression Feedback model but not the Retention model (Figure 6C). We also note that, consistent with our prior observations at the 4-cell stage (Carvalho et al., 2009) we did not observe evidence of turnover of ring myosin due to exchange with myosin in the cytoplasm. If ring myosin were turning over due to exchange with cytoplasmic myosin, we would expect the curve for fluorescence in the ring after the bleach to approach the control curve and the difference between the two curves to decrease exponentially. Instead, the two curves remained parallel and the difference remained constant (Figure 6C). Imaging after bleaching of the entire division plane at the 4-cell stage yielded a very similar result (Figure 6 - Figure Supplement 1). We conclude that acceleration of the per-unit-length constriction rate during closure, a conserved feature of contractile rings, does not arise from retention of components in the around-the-ring direction. Our results are instead consistent with the idea that acceleration arises from positive feedback between ring myosin and compression-driven cortical flow along the axis perpendicular to the ring.

**Figure 6.**
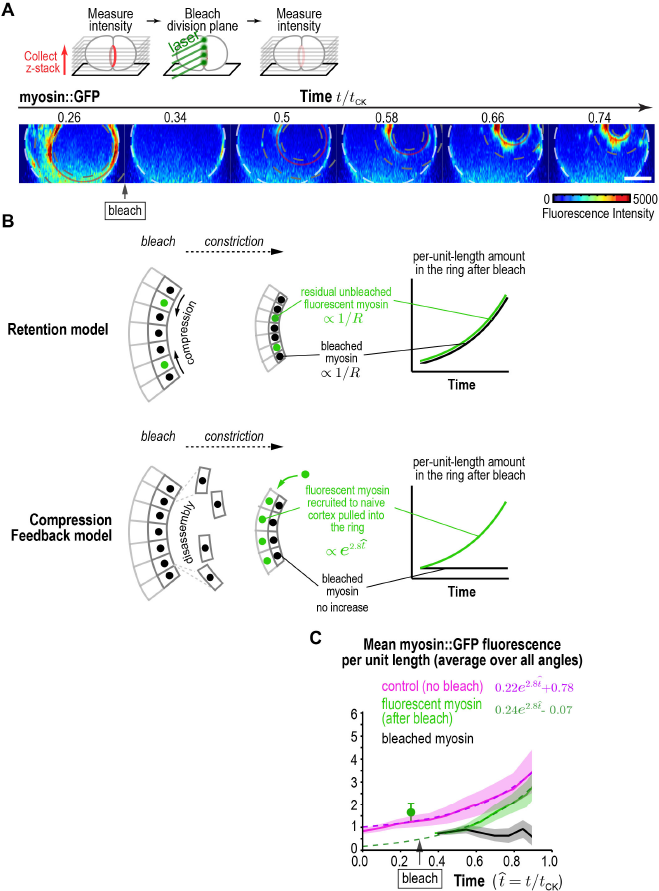
Fluorescence recovery after photobleaching of the division plane supports the Compression Feedback model. (**A**) *(top)* Schematic of the photobleaching experiment. *(bottom)* Images of the division plane reconstructed from 30×1μm z-stacks of an embryo expressing myosin::GFP whose division plane was bleached at t/t_CK_ ~0.3. Red circle marks the contractile ring and dashed circles mark the boundaries used for ring intensity measurements. Image series is representative of 8 imaged embryos. (**B**) Schematics illustrate the expected results predicted by the Retention and Compression Feedback models. (**C**) Graph plotting the mean per-unit-length amounts of fluorescent myosin::GFP in the ring for control embryos *(pink,* n=24 embryos) and embryos in which the division plane was bleached at the indicated time *(green,* n=8 embryos). The amount of bleached myosin::GFP in the ring *(black)* was calculated as the difference between the control and after bleach curves. Solid continuous lines are the average curves with errors shown as shaded regions. Dashed lines are exponential fits to the data. Errors for the control and after bleach data are SD and errors for the difference are SEM. Scale bar is 10 μm.

## DISCUSSION

### The Compression Feedback model: a new explanation for the acceleration in the per-unit-length constriction rate during constriction

Our simultaneous analysis of cortical and contractile ring dynamics suggests a new explanation for the acceleration in the per-unit-length constriction rate that allows contractile rings to maintain a high closure rate despite their progressively decreasing perimeter (Biron et al., 2004; Bourdages et al., 2014; Calvert et al., 2011; Carvalho et al., 2009; Ma et al., 2012; Mabuchi, 1994; Pelham and Chang, 2002; Zumdieck et al., 2007). Rather than arising from an increase in the per-unit-length amount ring myosin due to retention, we propose that acceleration arises from an exponential increase in the per-unit-length amount ring myosin due to feedback between ring myosin and compression-driven cortical flow along the direction perpendicular to the ring (Figure 7). In our model, polar relaxation allows ring myosin to compress cortical surface along the pole-to-pole axis perpendicular to the ring, thereby increasing the amount of ring myosin. An increase in the per-unit-length amount of ring myosin, in turn, would lead to increased cortical compression, resulting in a feedback loop that drives an exponential increase in the per-unit-length amount of ring myosin. In this model, the overall amounts of myosin, anillin (and presumably other components) in the ring would remain relatively constant as the ring constricts (Figure 5 - Figure Supplement 1) due to a balance between loss due to disassembly-coupled ring shortening and accumulation due to feedback in the perpendicular-to-the-ring direction. Thus, the relatively constant overall levels would mask a dramatic restructuring of the ring during closure. We note that the model we propose here is reminiscent of early conceptual models of cytokinesis, which hypothesized that polar relaxation coupled to a global upregulation of surface tension could trigger a flow of tension-generating elements towards the equator that would compress into a circular band and initiate a feedback loop (Greenspan, 1978; Swann and Mitchison, 1958; Taber, 1995; White and Borisy, 1983; Wolpert, 1960; Zinemanas and Nir, 1987; Zinemanas and Nir, 1988).

**Figure 7.**
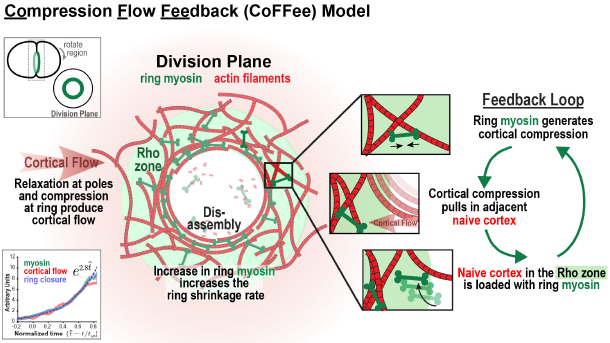
The Compression Feedback model: a feedback loop operating in the perpendicular-to-the-ring direction accelerates the per-unit-length constriction rate during ring closure. Schematic summary of the Compression Feedback model for cytokinesis. Polar relaxation allows ring myosin to compress cortical surface along the axis perpendicular-to-the-ring, which pulls more cortical surface that is loaded with myosin into the ring. Feedback between ring myosin and compression-driven cortical flow leads to an exponential increase in the per-unit-length amount of ring myosin that maintains the high overall closure rate as ring perimeter decreases.

In addition to ensuring timely cell content partitioning, an advantage of the feedback-based mechanism that we propose here is that it would render the ring robust to internal or external mechanical challenges, such as cell-cell contacts, obstacles in the crowded cell interior, or defects in the cytokinesis machinery. In all of these cases, a feedback loop between ring myosin and compression-based flow along the direction perpendicular to constriction would lead to the progressive build-up of contractile ring components until they reached a level where the obstacle could be overcome and constriction would again be able to proceed. Concentrating components by compression in the around-the-ring direction would not have this property, since successful constriction would be required to increase component levels. We note that similar ring-directed cortical flows have also been observed in the context of wound healing (sMandato and Bement, 2003), where they could potentially serve a similar function in allowing the cell to ramp up contractile force and achieve wound closure.

The experimental basis for our model is our analysis of cortical dynamics, which indicates that the compression of cortical surface within the ring along the axis between the relaxing poles that initiates during contractile ring assembly (Figure 3; (Reymann et al., 2016)), persists throughout constriction, resulting in a continuous flow of cortical surface into the ring. A second key finding is that the per-unit-length amount of ring myosin and anillin and the per-unit-length rates of cortical compression and ring constriction increase with the same exponential kinetics, suggesting control by positive feedback. We note that it remains possible that there is a distinct source of positive feedback (other than between ring myosin and cortical compression as we propose) that controls myosin recruitment, and that myosin levels in turn control the rates of constriction and cortical compression. However, since our data indicate that cortical surface is compressed within the ring, such a model would need to invoke an as yet uncharacterized process to explain why compression of the cortex within the ring would not increase the concentration of ring components. We note that compression within the ring along the direction perpendicular to the ring is also consistent with work in S. *pombe,* which has shown that contractile ring assembly occurs via a similar acto-myosin based compression of an equatorial band of nodes into a compact ring along the long axis of the cell (Vavylonis et al., 2008; Wu et al., 2006). However, in contrast to *pombe* where ring assembly and constriction occur in distinct phases, our model predicts that in animal cells, the accumulation of ring components due to compression along the direction perpendicular to the ring is ongoing, and serves to accelerate the per-unit-length constriction rate as the ring closes.

### Polar relaxation enables cortical compression within the ring along the pole-to-pole axis

Monitoring cortical dynamics in combination with laser ablation experiments revealed that the polar cortex is distinct from the cortex in the region between the contractile ring and the poles. The polar cortex expands in response to tension generated by the constricting ring, whereas the intervening cortex flows towards the ring without expanding. One possibility is that polar cortex is less stiff than the rest of the cortex, causing it to stretch and thin in response to ring constriction-induced tension. Alternatively, the polar cortex could turnover more rapidly, leading to a higher rate of surface renewal after stretching. A third possibility is that the polar cortex is more prone to rupture, repair of which would locally increase cortical surface. Consistent with this last idea, blebs have been reported at the cell poles in cultured vertebrate and *Drosophila* cells, where they have been proposed to release tension at the poles (Hickson et al., 2006; Sedzinski et al., 2011). The distinct mechanical properties of the polar cortex suggest that its composition could be different from that of the adjacent cortex. This idea is consistent with both older work suggesting the existence of mechanisms that clear contractile ring proteins from the poles (Bement et al., 2005; Chen et al., 2008; Foe and von Dassow, 2008; Murthy and Wadsworth, 2008; von Dassow, 2009; Werner et al., 2007; Zanin et al., 2013) and recent studies that have begun to uncover molecular mechanisms that may drive clearing. Work in *C. elegans* has demonstrated the existence of a mechanism in which Aurora A, localized to astral microtubules by association with its activator TPXL-1, actively clears contractile ring proteins from the polar cortex (Mangal et al., 2018). A reduction in f-actin intensity at the cell poles due to delivery of a phosphatase by segregating chromosomes has also been reported in *Drosophila* cells (Rodrigues et al., 2015). Understanding how the polar cortex is different in molecular and mechanical terms, and the mechanisms that generate these differences are important goals for future work.

Cleaving sea urchin embryos exhibit constriction kinetics essentially identical to those during the first division of the *C. elegans* embryo (Mabuchi, 1994). Pioneering work measuring the distance between surface-adhered particles and the behavior of pigmented cortex-associated granules (Dan, 1954; Dan and Dan, 1940; Dan et al., 1938), indicated that sea urchin embryos also exhibit a similar pattern of cortical expansion during ring constriction, in this case, a wave of cortical expansion that initiates at the poles and propagates through to the region adjacent to the furrow (Dan et al., 1938; Dan and Ono, 1954; Dan et al., 1937; Gudejko et al., 2012; Swann and Mitchison, 1958). Cortical compression and expansion have not been mapped in vertebrate cells; however, monitoring of fluorescent latex spheres adhered to cell surface proteins (Fishkind et al., 1996; Wang et al., 1994), injected stabilized fluorescent actin filaments (Cao and Wang, 1990), and fluorescently labeled myosin II (DeBiasio et al., 1996) all revealed concerted cortical flow towards the division plane in the equatorial region of the cell that contrasted with random surface movements at the cell poles. These observations suggest that feedback in which relaxation enables compression-driven cortical flow may be a conserved feature of animal cell cytokinesis.

### The Compression Feedback model predicts that the evolution of component levels in the ring during constriction requires both de novo recruitment and compression-driven cortical flow

It is worth noting that our proposed model represents an interesting twist on an ongoing debate in the cytokinesis field as to whether contractile ring components are delivered into the ring via cortical flow (Cao and Wang, 1990; DeBiasio et al., 1996; Fishkind et al., 1996; Wang et al., 1994) or recruited de novo from the cytoplasm downstream of RhoA-based signaling (Vale et al., 2009; Yumura, 2001; Zhou and Wang, 2008). In the Compression Feedback model, we propose that following anaphase onset contractile ring components are initially recruited to the equatorial cortex de novo, as has been observed (Vale et al., 2009; Yumura, 2001; Zhou and Wang, 2008), but then component levels are amplified by a feedback loop in which compression of cortical surface in the ring pulls new cortex into the Rho zone that is then loaded de novo with contractile ring components. Thus, during the exponential increase in the per-unit-length amount of ring components, compression-driven flow of new cortex into the Rho zone would be required for the subsequent de novo loading of contractile ring components. We would therefore propose that both the de novo loading of components by Rho-based signaling and compression-driven flow could contribute to the evolution of the component levels in the ring during constriction.

### The Compression Feedback model as a tool to describe the feedback-mediated evolution of the contractile ring

To quantitatively explore the idea that a feedback loop between the amount of ring myosin and compression-driven flow of cortical surface into the ring drives component accumulation during constriction, we developed an analytical mathematical framework, which we call the Compression Feedback model. The Compression Feedback model consists of three equations with three model parameters that describes this feedback and can recapitulate our experimental results. In addition to describing the processes underlying the evolution of the contractile ring, the Compression Feedback model provides a simple framework that can be used to analyze the consequences of molecular perturbations. Since the Compression Feedback model accurately describes the dynamics of the contractile ring and associated cortical network, an additional interesting future direction will be to use parameter changes derived from the Compression Feedback model as input for a finite-element model (similar to (Turlier et al., 2014)) in order to predict the evolution of cell shape given an *a priori* knowledge of cortical and contractile ring dynamics.

## METHODS

### C. elegans strains used in this study

| Strain Name | Genotype | Reference |
| --- | --- | --- |
| OD821 | *ItSi200[pOD1997; Pnmy-2::nmy-2::gfp; cb-unc-119(+)] II; unc-119(ed3) III* | This study |
| OD857 | *ItSi200[pOD1997; Pnmy-2::nmy-2::gfp; cb-unc-119(+)] II; unc-119(ed3); ruls32[pAZ132; pie-1/GFP::histone H2B] III* | This study |
| OD858 | *ItSi803[pOD1998; Parx-7::GFP::arx-7; cb-unc-119(+)] II; unc-119(ed3) III;* | This study |
| LP162 | *nmy-2(cp13[nmy-2::gfp + LoxP]) I* | (Dickinson et al., 2013) |
| OD95 | *unc-119(ed3) III; ItIs37 [pAA64; Ppie-1::mCherry::his-58; unc-119(+)] IV; ItIs38 [pAA1; Ppie-1::GFP::PH(PLC1delta1); unc-119 (+)]* | (Essex et al., 2009) |
| OD3011 | *ItSi1123[pSG017; Pani-1::GFP::ani-1 RE-encoded-exon5::ani-1 3'-UTR; cb unc-119(+)]II;unc-119(ed3)III* | This study |
| GOU2047 | *cas607[arx-2::gfp knock-in] V* | (Zhu et al., 2016) |

The *C. elegans* strains listed in the table were maintained at 20°C using standard methods. OD821 and OD858, expressing NMY-2::GFP, GFP::anillin, and GFP::ARX-7 were generated using a transposon-based strategy (MosSCI; (Frokjaer-Jensen et al., 2008)). Genomic regions encoding *nmy-2* (including 2079 bp and 1317 bp up and downstream of the stop codon, respectively), *ani-1* (including 2015 bp and 1215 bp up and downstream of the stop codon), and *arx-7* (including 3056 bp and 634 bp up and downstream of the stop codon) were cloned into pCFJ151 and sequences encoding GFP were inserted either just before *(nmy-2)* or after *(arx-7* and *ani-1)* the start codon. The single copy *nmy-2* transgene was generated by injecting a mixture of repairing plasmid (pOD1997, 50ng/μL), transposase plasmid (pJL43.1, Pglh-2::Mos2 transposase, 50ng/μL), and fluorescence selection markers (pGH8, Prab-3::mCherry neuronal, 10ng/μL; pCFJ90, Pmyo-2::mCherry pharyngeal, 2.5ng/μL; pCFJ104, Pmyo-3::mCherry body wall, 5ng/μL) into EG6429 (ttTi5605, Chr II). Single copy *ani-1* and *arx-7* transgenes were generated by injecting a mixture of repairing plasmid (pSG017 *(ani-1)* or pOD1998 *(arx-7),* 50ng/μL), transposase plasmid (CFJ601, Peft-3::Mos1 transposase, 50ng/μL), selection markers (same as for *nmy-2* strain) and an additional negative selection marker (pMA122; Phsp-16.41::peel-1, 10ng/μL) into EG6429 (ttTi5605, Chr II). After one week, progeny of injected worms were heat-shocked at 34°C for 2-4 hours to induce PEEL-1 expression and kill extra chromosomal array containing worms (Seidel et al., 2011). Moving worms without fluorescent markers were identified and transgene integration was confirmed in their progeny by PCR spanning both homology regions in all strains.

### C. elegans *RNA-mediated interference*

Double stranded RNA (dsRNA) targeting *arx-2* (K07C5.1) at a concentration of 1.7 mg/ml was generated by synthesizing single-stranded RNAs in 50μL T3 and T7 reactions (MEGAscript, Invitrogen, Carlsbad, CA) using cleaned DNA template generated by PCR from N2 DNA using the oligos (TAATACGACTCACTATAGGTCAGCTTCGTCAAATGCTTG and AATTAACCCTCACTAAAGGTGCAATACGCGATCCAAATA). Reactions were cleaned using the MEGAclear kit (Invitrogen, Carlsbad, CA), and the 50 μL T3 and T7 reactions were mixed with 50μL of 3x soaking buffer (32.7mM Na_2_HPO_4_, 16.5mM KH_2_PO_4_, 6.3mM NaCl, 14.1mM NH_4_Cl), denatured at 68°C for 10min, and then annealed at 37°C for 30 min to generate dsRNA. L4 hermaphrodite worms were injected with dsRNA and allowed to recover at 16°C for 44-50 hours prior to imaging.

### Generating a 4D map of cortical flow

Cortical flow was monitored in embryos expressing myosin::GFP obtained from adult hermaphrodites by dissection. Embryos were mounted followed by sealing with a coverslip on double thick (1 mm) low percentage agarose (0.5%) pads to prevent compression that biases the initial angle of furrow ingression (Figure 1 - Figure Supplement 1B). Images were acquired on an inverted microscope (Axio Observer.Z1; Carl Zeiss) equipped with a spinning-disk confocal head (CSU-X1; Yokogawa) and a 63x 1.40 NA Plan Apochromat lens (Zeiss) using a Hamamatsu Orca-ER digital camera (Model C4742-95-12ERG, Hamamatsu photonics). Images were collected using custom software, written in Python, that utilizes the Micro-Manager (open source software, (Edelstein et al., 2014)) microscope control library. A 3 x 0.75 μm z-series was collected (400ms exposure, 10-20% laser power) every 2s. After 15 time points, a 15 x 1μm z-stack, offset by 3μm from the cortical surface, was imaged to monitor the position of the closing contractile ring. The entire imaging series was repeated every 36s until the end of cytokinesis. Cortical flow was measured in maximum intensity projections of the 3 x 0.75μm z-stacks of the cortical surface, after orientation of the images to place the embryo anterior at the top and the posterior at the bottom, by correlating myosin fluorescence between consecutive images using Gunnar Farnebäck’s algorithm (Farnebäck, 2003) implemented within the openCV library with a 30-pixel window size. The threshold was calculated for every image by maximizing the ratio of total intensity inside a 200×350 pixel box positioned in the center of the embryo to the total intensity outside that box.

### Measurement of contractile ring position and size

Automated methods were employed to identify the edges of the embryo, determine the position of the contractile ring, and reconstruct the rings for each time point in an end-on view to determine the initial ingression axis (Figure 1 - Figure Supplement 2). Ring size and position were determined using custom Python software that: (1) identifies the orientation of the anterior-posterior (AP) axis and rotates the embryo to place the embryo anterior at the top and the embryo posterior at the bottom, (2) finds the embryo center in different x-z planes along the AP axis and calculates embryo radius, and (3) calculates the radius of the contractile ring and determines its position within the division plane. Details of each step are outlined below.

<u>Orienting embryos with their anterior end to the top:</u> Acquired z-plane images were convolved with a 10-pixel Gaussian kernel to reduce noise. An optimal signal threshold that partitioned the embryo interior from exterior was identified by finding a local minimum in the intensity histogram that produced a binary mask with expected area (~120000±50000 pixel^2^). The orientation of the AP axis was identified by fitting an ellipse to the thresholded area in the middle plane of the z-stack. The anterior side was identified by higher cortical myosin fluorescence and all images were rotated to place the embryo anterior at the top of the image and the embryo posterior at the bottom.

<u>Defining the central axis of embryo and determining embryo width:</u> The central axis of the embryo was defined by drawing a horizontal line across the oriented embryo at the midpoint between its anterior and posterior ends and identifying the first and last points along this line with signal above the threshold for each z-plane. The identified pixels were virtually projected in an end-on (x-z) view and fit to a circle by minimizing residuals. To account for fluctuations in the embryo boundary due to noise and fluorescence variation, the procedure was repeated 9 more times after shifting the position of the horizontal line towards the anterior pole by 10 pixels, covering approximately 1/5 of the embryo length (500 pixels). The position of the AP axis and the radius of the embryo were determined by averaging the 10 measurements.

<u>Measuring contractile ring size and position:</u> As illustrated for the central plane images shown in Figure 1 - Figure Supplement 2, the position of the contractile ring was determined by identifying pairs of points with the highest myosin fluorescence intensity on the opposite edges of the embryo in each z-plane that were not more than 20 pixels apart in the horizontal direction and were located at a y-axis position near the embryo middle. Contractile ring radius and position were determined by projecting the points to generate an end-on (x-z) view and fitting the data with a circle. The ring fit was iteratively improved by calculating predicted positions of myosin fluorescence at the ring in each z-plane using initially fitted parameters. Intensity maxima within 5 pixels of the predicted location were identified and the ring was refit. The initial guesses for the contractile ring size and position at the next time point were estimated from the previously calculated ring values. The algorithm restricted ring position fluctuations to 20 pixels along anterior-posterior axis and the size was estimated assuming constant rate of ring constriction. The automatic ring measurements were manually confirmed for each embryo. The initial ingression axis was determined as illustrated (Figure 1 - Figure Supplement 2) by fitting a line through the centers of the rings with a normalized ring size 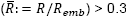.

### Embryo time alignment for averaging

Sequences from individual embryos were time aligned by defining zero time (t_0_) and the total time of cytokinesis (t_CK_) for each embryo, and normalizing time by *t_CK_* prior to averaging, 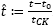. An initial determination of t_0_ and *t_CK_* was made by fitting a line to the plot of normalized ring size 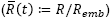 versus time between 30% and 80% closure for each embryo as outlined in Figure 1A. Extrapolation of this line for each embryo defined *t_0_* as the time where the fitted line intersects 1, and the time of cytokinesis, *t_CK_* as the time where the fitted line intersects 0. Due to the small number of measurements from each embryo available for fitting (3-5 values where 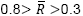), the values of t_0_ and *t_CK_* were refined by fitting 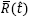 for each embryo to the average dimensionless ring size, 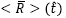. Calculation of the average dimensionless ring size was performed in iterative manner. The time for each embryo was aligned by t_0_ and normalized by *t_CK_* using estimates from the fitted line in the first iteration. The average dimensionless ring size 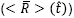 was calculated by averaging normalized ring sizes of all embryos at corresponding normalized time. Contractile ring size was approximated for intermediate time points by linear interpolation. In further iterations, t_0_ and *t_CK_* were refined for every embryo by minimizing the residuals between its normalized ring size, 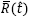, and the average dimensionless ring size, 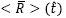, throughout the entire timecourse of cytokinesis, thus increasing the number of time points available for fitting t_0_ and *t_CK_* (6-10 values per embryo). After refining time alignment and normalization for each embryo, average dimensionless ring size was re-calculated and t_0_ and *t_CK_* were refined for each embryo again. The refinement process was repeated until changes in average dimensionless ring size, 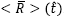, were smaller than 0.001 on average (achieved within a few iterations). The collective fitting of all t_0_ and *t_CK_* at every iteration was performed under restriction that the line fit through 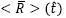 between 0.8 and 0.3 intercepted 0 at 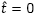 and 1 at 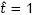. This restriction ensured that t_0_ and *t_CK_* determined from fits of individual embryos to the average ring size would be consistent with their original definition. The dimensional ring kinetics, < *R >* (t), can be recovered using the following equation

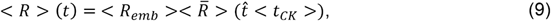

where < *R_emb_* >= 14.7 ± 0.7 *μm* and < *t_CK_* >= 200 ± 30 *s* are average embryo radius and time of cytokinesis accordingly.

### Cortical flow averaging

Cortical flow averaging was performed after spatial and temporal alignment of data collected in different embryos (n=93 embryos from 93 worms filmed over the course of 5 days for control, Video 2; n=68 embryos from 68 worms filmed over the course of 4 days for *arx-2(RNAi),* Video 3). The number of embryos was chosen to achieve at least 10-fold coverage for all areas of the cortical map for controls and 5-fold coverage for *arx-2(RNAi).* Linear interpolation was used to approximate the flow between consecutive time points. Because our imaging regime required periodic z-stack acquisition to determine the trajectory of ring closure, no flow approximation was done during those time periods (~6s gap every 30s). The flow data for each time point was represented as a set of vectors with direction and magnitude corresponding to the direction and magnitude of the cortical flow at the base of the vector. The base of each vector had two spatial coordinates: *x,* the position along the anterior-posterior axis (where the position of the contractile ring was defined as 0), and θ, the angular position relative to the initial ingression axis (defined as described in Figure 1A and Figure 1 - Figure Supplement 2). We note that mitotic exit is accompanied by a brief (~50-60s) period of rotational flow ((Naganathan et al., 2014; Schonegg et al., 2014); see Video 1), which dissipates soon after initiation of cytokinesis 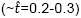. As this rotational contribution is not relevant here, we removed it by averaging the data from the right and left halves of the embryo (in an end-on view), allowing us to focus on rotation-independent flows. Thus, the flow with angular positions greater than 180 degrees was mirrored in angular direction

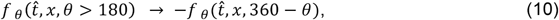

*f _θ_* is the angular component of the flow vector 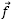. The flows were normalized by the embryo size and cytokinesis rate 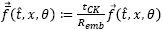 and averaged according to its position and time

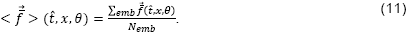

### Calculation of expected cortical surface flow profiles

To aid in the interpretation of experimental results, expected profiles for cortical surface movement were calculated for defined patterns of cortical surface increase and plotted (Figure 1B and Figure 1 - Figure Supplement 3). The general form of surface movement velocity is given by the following equation

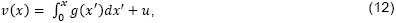

where *g(x)* is the amount of cortical surface gain and *u* is the velocity of asymmetric ring movement, which could be positive or negative, depending on whether the ring is moving towards or away from the surface. From equation (12) we obtain the following predictions

Uniform surface increase: *v*(*x*) = *Cx* + *u*;
Polar surface increase: *v*(*x*) = *C + u*;
Behind the ring surface increase: *v(x)* = *u* (if the asymmetry of cytokinetic furrowing arises due to global surface movement) or *v*(*x*) = 0 (if the asymmetry in surface increase is related to the asymmetric furrowing).

### Cortical laser ablation

Cortical laser ablations, presented in Figure 2, were performed using a robotic laser microscope system (RoboLase) (Botvinick and Berns, 2005). Embryos expressing myosin::GFP were mounted using standard procedures. A cortical cut, approximately 10 μm long, was made on the anterior side of the embryo when the ring was at ~50% closure (7μm radius). The cut was confirmed by comparison of cortical fluorescence images before and after the cut and was considered successful if the foci moved away from the cut area (~3.5μm distance), indicating cortical tension release. Contractile ring closure rate was calculated by measuring the difference in ring sizes before and after the cut, assessed from two 4×2μm z-stacks acquired immediately before the cut and 13s later. Errors in measuring the radius at the two timepoints were determined from the procedure used to fit the data to a circle and were propagated to determine the errors in the constriction rate measurements for individual embryos; mean errors are S.E.M. The cortical opening after ablation was approximately 35μm^2^; this translates into an additional reduction in ring radius by ~0.8μm, if the cortical surface tension dominates the ring closure rate. This additional decrease in ring size within 13s should correspond to increase of the control rate (0.22μm/s) by ~30% (0.06μm/s). The experiment was repeated 19 times for no cut condition, 14 times for parallel cut, and 15 times for perpendicular cut. All imaging was performed over the course of 5 days. The number of embryos was chosen to achieve sufficient accuracy in the determination of mean ring closure rates to assess whether it was altered by the cuts.

### Calculation of the surface area flowing into the division plane

We calculated the amount of surface area flowing into the division plane from flow measurements made 7 μm away from the position of the furrow on the anterior and posterior sides (as illustrated in Figure 3B). The rate of the surface flow is

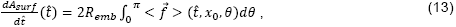

where *x_0_* is -7 μm and 7 μm for the rate of flow from the anterior or the posterior sides, respectively. The total amount of surface area that entered the division plane from any time ř_0_ to ř is obtained by integrating equation (13) over time

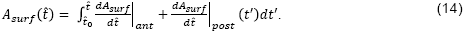

The increase in area of the division plane was calculated as following

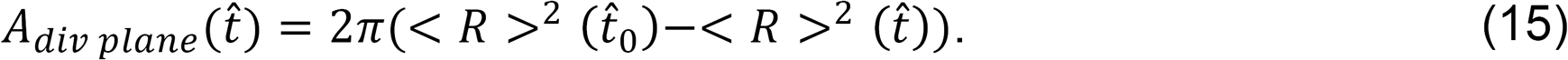

In Figure 3B we used 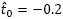. The cortical surface area compressed in the ring can be inferred from the difference between the surface area entering the division plane and the area of the division plane

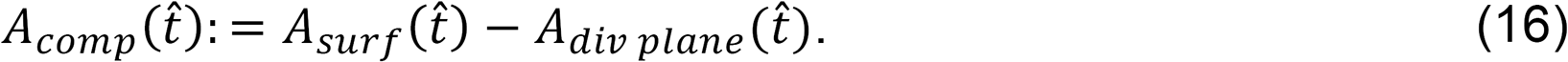

### Division plane imaging

For quantification of myosin::GFP and GFP::anillin amounts in the contractile ring, adult worm dissection and one-cell stage embryos imaging was performed in a custom microdevice (Carvalho et al., 2011). The device was mounted on an inverted microscope (Axio Observer.Z1; Carl Zeiss) and embryos were imaged with a 63×1.4NA Plan Apochromat objective using an electron-multiplying charge-coupled device camera (QuantEM:512SC, Photometrics; 100ms exposure, EM gain set to 500, 10% laser power). Division planes were reconstructed from 40 x 0.5μm z-stacks collected every 30s after background subtraction and attenuation correction. All imaging was done at 20°C.

### Contractile ring photo-bleaching and imaging

1-cell stage embryos were mounted in microdevices as for division plane imaging and 4-cell stage embryos were mounted on slides with 2% agarose pads. Embryos were imaged on a Nikon TE2000-E inverted microscope equipped with a 60×1.40NA objective, an EM-CCD camera (iXon; Andor Technology; EM-Gain=220, Exposure =100ms), and a krypton-argon 2.5 W water-cooled laser. For 1-cell stage embryos, division planes were reconstructed from 30×1 μm stacks acquired every 20s with 20% laser power and photo-bleaching was performed by 2 sweeps of a 488nm laser with 100% power and 500μs dwell time. For 4-cell stage embryos, division planes were reconstructed from 16×1 μm stacks acquired every 10s with 50% laser power and photo-bleaching was performed by 2 sweeps of a 488nm laser with 100% power and 100μs dwell time. For 4-cell stage embryos, the time between the prebleached and first postbleached images was 6s.

### Estimation of depth attenuation

To estimate depth attenuation within the division plane, we quantified the intensity of the division plane in two cell embryos expressing a GFP-tagged probe expected to be uniformly present on the plasma membrane. From each image, we subtracted a background intensity calculated as the average value inside two 11×11 μm rectangles positioned 2 μm away from the division plane inside the anterior and posterior cells (Figure 3 - Figure Supplement 3). The division plane intensity profile was obtained by performing a 30 pixel maximum intensity projection along the AP axis, with the division plane positioned approximately in the middle (Figure 3 - Figure Supplement 3). The intensity profiles in z from 13 embryos were fitted to an exponential using the same characteristic attenuation depth for all embryos

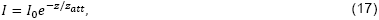

which yielded a characteristic depth of attenuation, *z*_att_, of 15 μm.

### Quantification of myosin and anillin intensity in the contractile ring and on the cortex

For embryos at the 1-cell stage, myosin::GFP and GFP::anillin intensities in the contractile ring and on the cortex were quantified in 40×0.5 μm z-stacks containing the ring after correction for depth attenuation and subtraction of background fluorescence. Average intensity along the ring was calculated across a set of embryos in 30 degree arcs (for myosin::GFP, n=36 embryos from 18 worms filmed over 5 days; for anillin::GFP, n= 26 embryos from 14 worms filmed over 4 days). The number of embryos was chosen to determine mean fluorescence with sufficient accuracy to derive appropriate conclusions. Positions along the ring were referenced based on the angle between the line from the position on the ring to the ring center and the initial ingression axis. Linear interpolation in time was used for every embryo to estimate intensity in the intermediate time points to perform averaging. Measured intensities were divided by arc length and averaged between different embryos to obtain mean GFP fluorescence per-unit-length for different angular ranges and the average for all angles. Total ring GFP fluorescence was calculated by integrating over ring perimeter. Cortical intensities were quantified by choosing the time point with the ring size closest to 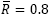 and measuring total fluorescence in the 15^th^ plane after correction for depth attenuation and subtraction of background fluorescence.

Measurements of myosin::GFP fluorescence in the ring at the 4-cell stage were performed as described in Carvalho et. al., 2009. However background fluorescence was determined as the mean fluorescence within a variable size circle at least 10 pixels in diameter, instead of fixed at 10 pixels, to improve measurement quality.

### Derivation of the Compression Feedback model for cytokinesis

The Compression Feedback model formalizes the following conceptual view of cytokinesis: After anaphase onset spindle based signaling patterns the cortex, generating an equatorial zone where RhoA promotes the recruitment of contractile ring components (the Rho zone). Within the Rho zone, myosin engages with actin to exert an isotropic force that compresses the cortical surface, resulting in uniform compression across this region, as is observed experimentally (Figure 3A). Due to polar relaxation, the compressing cortex pulls näive cortex not previously patterned by RhoA signaling into the Rho zone. We propose that the new cortical surface that flows into the Rho zone as a result of compression is also loaded with contractile ring components. Thus, a feedback loop is established along the direction perpendicular to the ring, in which myosin in the ring compresses cortical surface, which pulls more surface that is loaded with myosin into the ring. Disassembly in the around-the-ring direction reduces ring components in proportion to the reduction in length, and does not alter the per-unit-length amount of myosin. Thus, changes in myosin levels are determined solely by the flow of näive cortex into the Rho zone along the direction perpendicular to the ring, which can be solved as a one-dimensional problem. We assume that the rate of compression of cortical surface (between *x* and *x + dx)* is proportional to local myosin concentration, *m(x,* t), which exerts stress onto the actin network resulting in

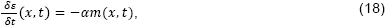

where *ε* is the cortical strain (i.e. change in length of cortical surface per-unit-length) and α is a proportionality constant that reflects the ability of the cortex to be compressed by ring myosin. The velocity of cortical surface movement is obtained from the following relationship (see also equation (12)).

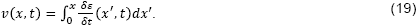

The conservation of mass for myosin flow results in the following

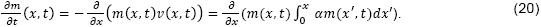

If we integrate equation (20) over x on *(-w, w)* domain we obtain

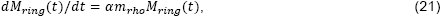

where 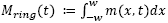 is the total per-unit-length amount of ring myosin engaged in compression, 2w is the width of the Rho zone/contractile ring where myosin is engaged and compressing cortex and *m_rh0_ ■■= m(w, t)* is the concentration of myosin loaded onto the cortex when it enters the rho zone. The velocity of flow of näive cortex into the rho zone is

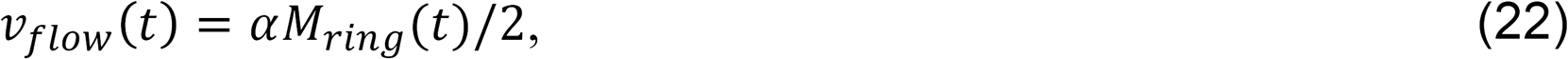

The one half is included to account for the fact that flow comes in from both sides. The solution of equation (21) is

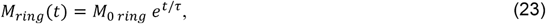

where we define the characteristic time of myosin accumulation, τ, as 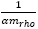. Note that the total amount of myosin in the ring will be the amount of engaged ring myosin plus an added baseline that would include any myosin not involved in compression (see equation Error! Reference source not found.). We assume the per-unit-length rate of ring shrinkage is proportional to the amount of ring myosin, as observed in our data,

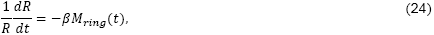

where *β* is a proportionality coefficient that reflects the ability of the ring to be constricted by ring myosin. Using equations (23) and (24), we obtain the dynamics of contractile ring size over time

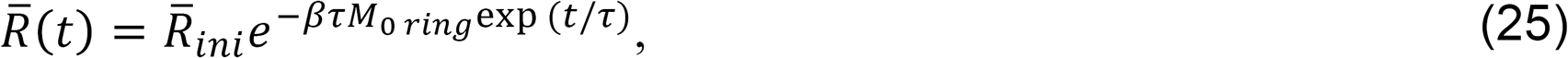

where 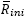 is the dimensionless characteristic size of the ring; essentially the radius at minus infinity if the same exponential process controlling contractile ring assembly extended back in time infinitely. Instead, *in vivo* cytokinesis initiates when spindle-based signaling activates RhoA on the equatorial cortex leading to the abrupt recruitment of contractile ring components. If the time frame of reference is chosen so that *t* = 0 is cytokinesis onset immediately following the initial patterning of the cortex by RhoA, *M_0 ring_* is the amount of ring myosin immediately following this event and the initial size of the ring is

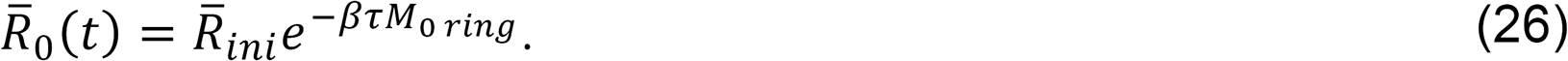

To facilitate future use of our model for analysis of contractile ring closure data, we use the time frame of reference where *t* = 0 is the point of 50% closure (i.e. 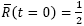), an easily identifiable time point that does not rely on exact assessment of the precise onset of cytokinesis. In this reference, 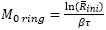 and by defining dimensionless velocity as 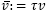, we obtain equations (4-8). Note that equation (4) can be rewritten in the following way

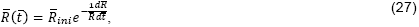

where 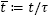. This relationship implies that in this dimensionless time, where *t =* 0) = !, any two rings of the same size have the same dimensionless constriction rate.

### Data availability

All data is available from the authors upon request.

### Code availability

The custom computer code used in this study is freely available from: https://github.com/renatkh/cytokinesis.

## ACKNOWLEDGEMENTS

This work was supported by a fellowship from the Jane Coffin Childs Memorial Fund to R.N.K. and grants to M.W.B from AFOSR (FA9550-08-1-0284) and the Beckman Laser Institute Foundation. J.S.G-C was supported by the University of California, San Diego Cancer Cell Biology Training Program (T32 CA067754). A.D. and K.O. receive salary and other support from the Ludwig Institute for Cancer Research. We would also like to thank Michael Glotzer for discussions that helped us align our model with current thinking about the Rho zone.

---

**Figure 1 - Figure Supplement 1.**
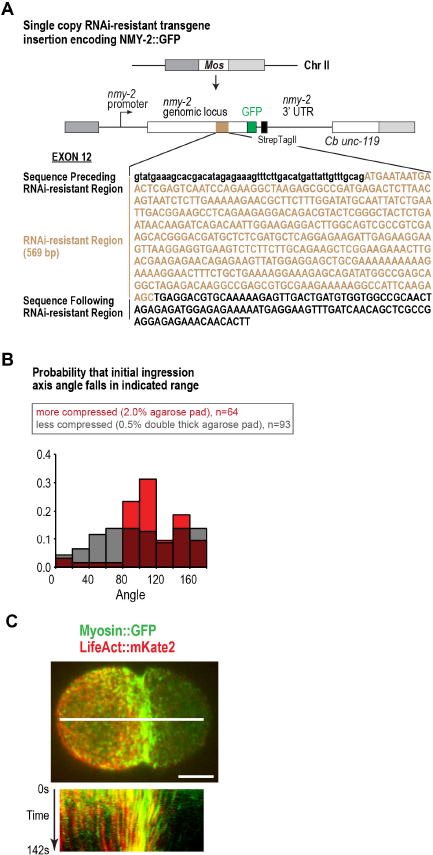
Actin and myosin move together with the cortical surface during cytokinesis. (**A**) Schematic of the single-copy *nmy-2::gfp* transgene inserted into a specific locus on chromosome II. Cb *unc-119*, the *unc-119* coding region from the related nematode *C. briggsae,* was used as a transformation marker. The transgene was re-encoded while maintaining amino acid sequence in the indicated region to render it resistant to RNAi targeting the endogenous gene for other experiments, this feature was not used in the experiments in this manuscript. **(B)** Compression biases the direction of contractile ring closure. Graph plotting the probability that the angle between the objective axis and the initial ingression axis falls in the indicated range for embryos mounted with more *(red)* or less *(grey)* compression. Due to this bias, embryos were mounted using the low compression conditions shown in grey. (C) Actin and myosin move together with the cortical surface during cytokinesis. The white line in the center of the image *(top)* indicates the region used for the kymograph *(bottom).* Image is representative of 5 imaged embryos. Scale bar is 10μm.

**Figure 1 - Figure Supplement 2.**
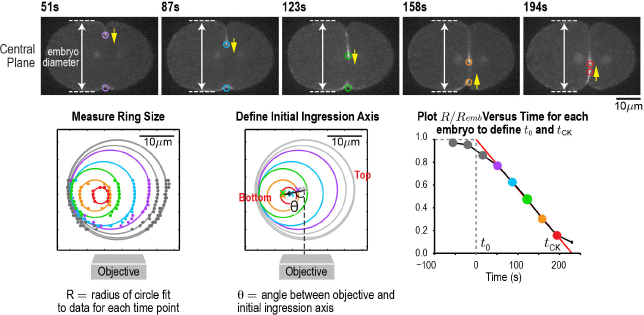
An automated method for monitoring contractile ring closure. *(top)* Central plane images of the embryo in Figure 1A. Panels on the lower left and lower right are reproduced from Figure 1A for comparison. An automated algorithm was used to identify the edges of the embryo (dashed lines) and the position of the contractile ring (colored circles) in each z-plane. Yellow arrows mark the direction of furrow ingression and illustrate how the furrow initially ingresses from the top and then changes directions to ingress from the bottom during the second half of cytokinesis. *(lower left)* Points marking contractile ring position in the z-planes were projected onto an end-on view of the division plane. Data for different timepoints in this representative embryo are shown in colors corresponding to the circles in the central plane images. Ring sizes were measured by fitting circles to the data. *(middle)* The initial axis of contractile ring closure was defined by the angle θ between the objective axis and a line fit through the centers of the contractile rings with a normalized size > 0.3. *(right)* = A plot of normalized ring size versus time for this embryo defines t_0_ and t_CK_ as the times when a line fit through the points corresponding to ring sizes between 0.3 and 0.8 crossed 1 and 0, respectively. Scale bar is 10μm.

**Figure 1 - Figure Supplement 3.**
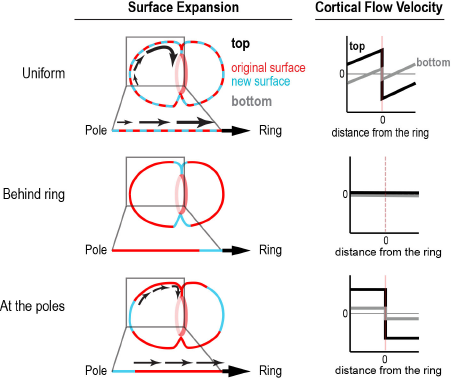
Different profiles of cortical surface velocity along the A-P axis are predicted for different spatial patterns of surface gain. *(top)* For surface gain behind the ring, no cortical movement is predicted on the embryo surface. *(middle)* For uniform surface gain, a gradient of velocities will be observed, where the cortex immediately behind the ring moves at the speed of the ingressing furrow, and cortical velocity decreases linearly towards the cell poles. *(bottom)* Reproduced from Figure 1C for comparison. If surface is gained only at the poles, cortical velocity will be constant in magnitude within the flow map region with opposite direction on the two sides of the ring.

**Figure 2 - Figure Supplement 1.**
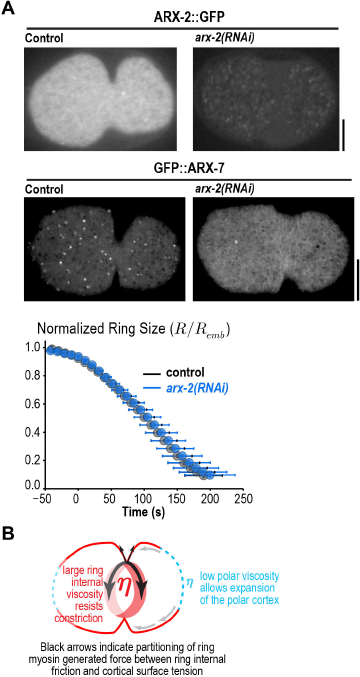
Arp2/3 depletion does not alter ring constriction kinetics. (**A**) Images of cortical ARX-2::GFP *(top)* and GFP::ARX-7 *(middle)* in control and *arx-2(RNAi)* embryos confirm loss of cortical Arp2/3 complex (images are representative of 10 imaged embryos for each condition in the GFP::ARX-7 strain and 15 for control and 13 for *arx-2(RNAi)* in the ARX-2::GFP strain). Scale bars are 10μm. *(bottom)* Graph plots average contractile ring size versus time for control (grey) and *arx-2(RNAi) (blue)* embryos expressing myosin::GFP (n= 93 embryos for control and 68 embryos for *arx-2(RNAi)).* Error bars are standard deviation. (**B**) Schematic illustrating the partitioning of ring myosin generated force between ring internal friction and cortical surface tension. Ring myosin generated force primarily counters ring internal friction to drive constriction. The low viscosity of the polar cortex causes it to expand when it comes under tension due to the constricting ring.

**Figure 3 - Figure Supplement 1.**
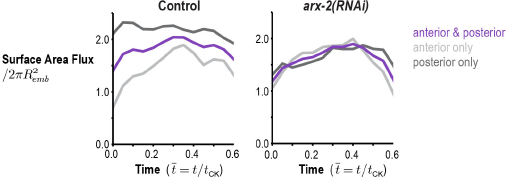
Arp2/3 inhibition abolishes the asymmetry in the amount of cortex entering the division plane from the anterior and posterior sides. Graphs plot the rate of cortical flux across the anterior *(light grey)* and posterior *(dark grey)* boundaries (see schematic in Figure 3B) versus the mean for the two sides *(purple)* for control and *arx-2(RNAi)* embryos. Calculated from the average flow maps for the control (n= 93 embryos) and *arx-2(RNAi)* (n= 68 embryos) conditions.

**Figure 3 - Figure Supplement 2.**
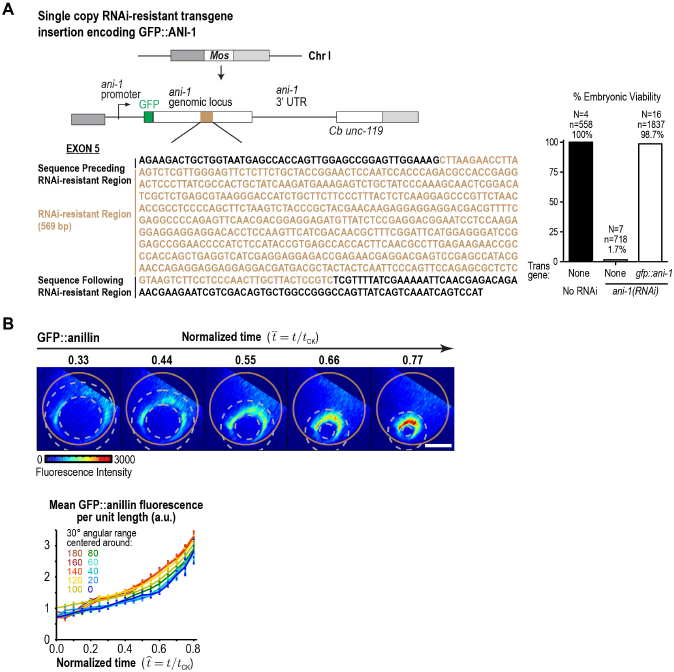
GFP::anillin fluorescence in the ring increases exponentially during constriction. (**A**) (*left*) Schematic of the single-copy *gfp::ani-1* trangene. The transgene was reencoded while maintaining amino acid sequence in the indicated region to render it resistant to RNAi targeting of the endogenous *ani-1* gene to allow testing of the functionality of the GFP::ANI-1 fusion. *(right)* Graph plotting embryonic lethality demonstrates that the *gfp::ani-1* transgene is functional. (**B**) *(top)* Images of the division plane in an embryo expressing GFP::anillin. *(bottom)* Graph plots GFP::anillin fluorescence per unit length of the ring for the indicated angular ranges. Error bars are the SEM.

**Figure 3 - Figure Supplement 3.**
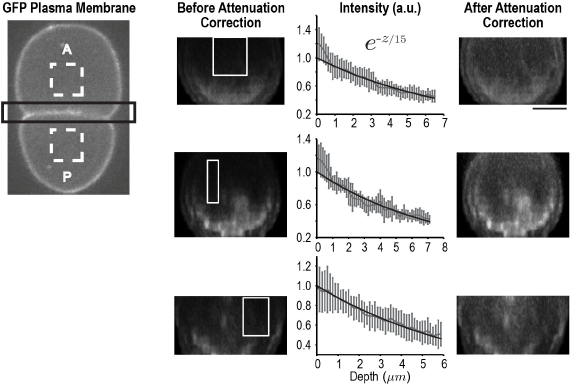
Correcting for signal attenuation with sample depth. Fluorescence attenuation with embryo depth was estimated from fluorescence intensity measurements made at the cell-cell boundary of the 2-cell embryos expressing a GFP-tagged plasma membrane marker. Cell-cell boundaries were reconstructed from 40 plane z-stacks. The intensity profile at each slice was calculated by subtracting the average background intensity estimated from dashed rectangles (left) from the cell-cell boundary region (black rectangle) at each slice and calculating the maximum intensity projection along AP axis. The effect of depth on signal was calculated from the reconstructed division planes by plotting the mean signal as a function of depth in 10 rectangular regions (white boxes) where the signal was expected to be uniform; three examples are shown here. All intensity profiles were simultaneously fitted using a single exponential. Error bars are the SD. On the right, the same cell-cell boundaries are shown after correction for depth attenuation. The scale bar is 10 μm.

**Figure 3 - Figure Supplement 4.**
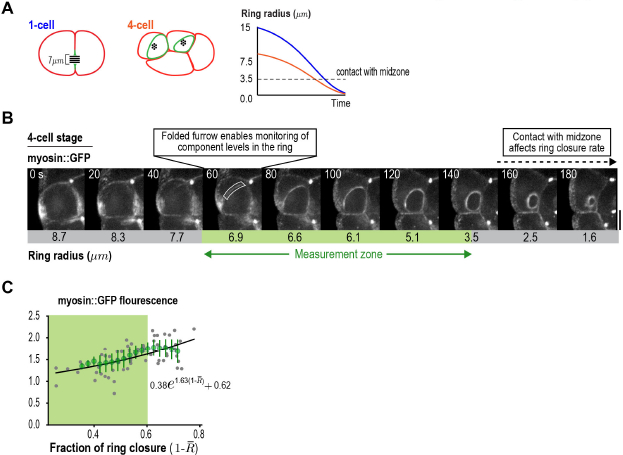
Ring component dynamics at the 4-cell stage are consistent with exponential accumulation. (**A**), *(left)* Schematic illustrating the relative geometries of cytokinesis in 1- and 4-cell stage *C, elegans* embryos. (right) The range of ring sizes between furrow formation and contact with the midzone, which occurs at a ring radius of about 3.5 μm in all divisions and alters constriction rate and component accumulation (Carvalho et al., 2009), is much smaller at the 4-cell stage than at the 1-cell stage. (**B**) Myosin levels in the ring can only be monitored over a limited range of ring size at the 4-cell stage. Images of the division plane in a representative dividing cell at the 4-cell stage reconstructed from 16×1pm z-stacks of an embryo expressing myosin::GFP (n=16 embryos imaged). The range of ring sizes for which myosin levels can be measured is indicated *(Measurement zone).* (**C**) Graph plotting measured mean per-unit-length myosin::GFP fluorescence in the ring at the 4-cell stage fit to an exponential equation with the same baseline contribution as the 1-cell stage data in Figure 3C *(blackline).* Error bars are the SEM.

**Figure 5 - Figure Supplement 1.**
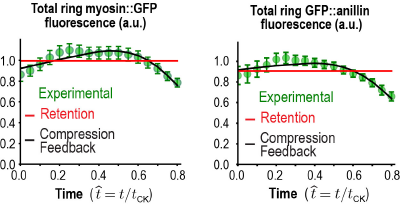
Total myosin::GFP and GFP::anillin in the ring. Graphs plotting mean total ring fluorescence (average over all angles; *green)* for myosin::GFP (n=36 embryos) and GFP::anillin (n=26 embryos). Error bars are the SEM. The predictions for the Compression Feedback *(black lines)* and Retention *(red lines)* models are also shown. Error bars are the SEM.

**Figure 6 - Figure Supplement 1.**
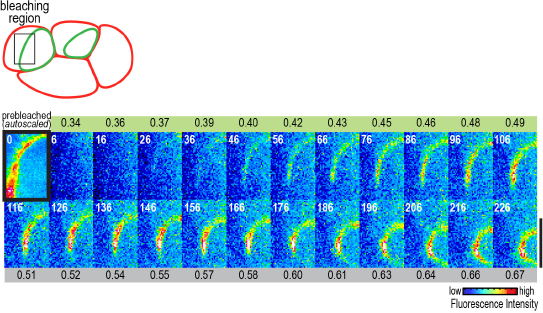
Recovery of myosin::GFP fluorescence after division plane bleaching at the 4-cell stage. To test whether compression-driven cortical flow delivers components to the ring at the 4-cell stage as well as at the 1-cell stage, we monitored recovery after photobleaching the entire contractile arc. Images show a representative bleached embryo (n=10). The observed recovery pattern was very similar to what we observed at the 1-cell stage. Scale bar is 10 μm.

## SUPPLEMENTARY VIDEO LEGENDS

**Video 1. Cortical flow imaged in a control embryo expressing myosin::GFP.**

Playback is 6x realtime. The video is constructed from maximum intensity projection of 3 x 0.75 μm plane z-stacks acquired at 2 s intervals. The red line marks the position of the division plane. The arrows represent the surface movement between consecutive frames at the base of the arrow. The length of the arrow is 5 times the magnitude of movement. The direction is also color coded according to the color wheel as shown in Figure 1B.

**Video 2. Average cortical flow map calculated from time lapse imaging of the cell surface in 93 control embryos expressing myosin::GFP.** *(top, left)* Schematic illustrates location of the cylindrical surface covered by the map. *(top, right)* Dynamic schematic illustrates ring size and position for each value of t/t_CK_. *(bottom, left)* The movement of each blue dot corresponds to surface movement at its location. The y-axis is the angular position relative to the initial ingression axis. The x-axis is the distance from the division plane along the anterior-posterior axis. *(bottom, right)* Dynamic graph plots the magnitude of the component of surface velocity aligned along the anterior-posterior axis for the top (150-180°; black) and bottom (0-30°; grey) regions of the cortex.

**Video 3. Average cortical flow map calculated from time lapse imaging of the cell surface in 68 *arx-2(RNAi)* embryos expressing Myosin::GFP.** *(top, left)* Schematic illustrates the location of the cylindrical surface covered by the map. *(top, right)* Dynamic schematic illustrates ring size and position for each value of t/t_ck_. *(bottom, left)* The movement of each blue dot corresponds to surface movement at its location. The y-axis is the angular position relative to the initial ingression axis. The x-axis is the distance from the division plane along the anterior-posterior axis. *(bottom, right)* Dynamic graph plots the magnitude of the component of surface velocity aligned along the anterior-posterior axis for the top (150-180°; black) and bottom (0-30°; grey) regions of the cortex.

